# Fixed points and multistability in monotone Boolean network models

**DOI:** 10.1101/2025.10.16.682751

**Authors:** Sarah Adigwe, BV Harshavardhan, Mohit Kumar Jolly, Tomáš Gedeon

## Abstract

Gene regulatory networks (GRN) control the expression levels of proteins in cells, and understanding their dynamics is key to potentially controlling disease processes. Steady states of GRNs are interpreted as cellular phenotypes, and the first step in understanding GRN dynamics is describing the collection of steady states the network can support in different conditions. We consider a collection of all monotone Boolean function models compatible with a given GRN, and ask which steady states are supported by most models. We find that for networks with no negative loops, there is an explicit hierarchy in the prevalence of individual steady states, as well as the prevalence of bistability and multistability. The key insight that we use is that monotone Boolean models supporting a given equilibrium are a product of prime ideals and prime filters of the lattices of monotone Boolean functions. To illustrate our result, we show that in the EMT network associated with cancer metastasis, the most common equilibria correspond to epithelial (E) and mesenchymal (M) states, and the bistability between them is the most common bistability among all network-compatible monotone Boolean models.

**Author summary:** Cells adjust their behavior in response to external inputs via networks of genes that regulate each other’s expression, until they arrive at a new steady state. Each interacting network of genes can behave in different ways that depend on internal and external cellular conditions. In this paper we consider, for a given network, an entire collection of particular type of models (monotone Boolean models) that represent all different ways that network can behave. Then, for any given state a network can potentially be in, we describe all monotone Boolean models that have that state as a steady state. We consider those states that are supported by more models to more likely represent the states that the network will be in. We apply our approach to EMT network that is important in cancer metastasis. We show that the most common steady states are those correspond to epithelial and mesenchymal states, and that the bistability between these two states is the most common bistability. This confirms the experimental results that these are the most common states of the EMT network.

## 1 Introduction

Unicellular organisms and cells in multicellular organisms solve complicated control problems related to the allocation of resources, division, and many others quickly and efficiently. Most of these tasks involve activation, deactivation, or expression of new proteins. The main conceptual model of how the chemical signal results in these changes takes the form of a *network* of interactions, where the presence of one signal results in increased or decreased activity/abundance of a downstream signal.

Many regulatory networks have been constructed [1–7] for organisms and tasks ranging from *Escherichia coli* to humans. An important question with obvious biological implications is to understand the range of dynamics a given network can support. In particular, a first step is to describe the steady states the network can support, as these are interpreted as different phenotypes that the network can support.

Experimentally, this is an untractable question, since the answer depends on both the external conditions and the internal state of the cell. Using mathematical models to answer this question is difficult, as the answer may depend on the choice of a model, its parameterization, and the choice of an initial condition. Unknown parameters can be addressed using parameter sampling methods [8] or using combinatorial parameterization of all ODE models with steep nonlinearities [9, 10], but both of these approaches quickly approach their limits when dealing with large networks. An alternative modeling approach of using a Boolean network model [11–13] has the (apparent) advantage of not needing any parameters, but the choice of update functions is a form of parameterization. Often, these models analyze a single, or only a few, Boolean models compatible with the network raising the question of whether some potential dynamics have been missed.

We propose a different approach [14] by constructing a finite collection of all monotone Boolean models (MBM) compatible with the network [15]. These are constructed as collections of monotone Boolean functions [16] (MBF) that update the state of each node based on its inputs. The monotonicity of the Boolean function reflects the network activation vs. deactivation edges. Monotone Boolean functions can be constructed by induction, but their size grows rapidly [17, 18]. In this approach, the network dynamics is a collection of all dynamics supported by all monotone Boolean models, which is weighted in importance by its prevalence. That is, if a particular steady state is supported by 80% of all MBMs, then it is more important than a steady state that is only supported by 10% of all MBMs.

Since the number of MBMs for a given network is finite, the prevalence of all dynamics is, in principle, computable. However, the number of MBM grows very quickly, and even computing steady states for a single MBM is NP-complete [19, 20].

In this paper, we concentrate on NL-free networks, which are networks that have no negative loops (cycles). We show that for such networks, the MBMs that support particular equilibria have a structure of products of down- and up-sets in lattices of MBFs. As a result of this characterization, we are able to completely describe the prevalence of each Boolean steady state, bistability, and multistability for any such network. This leads to a general computational procedure that we illustrate on monotone Boolean functions with *k* = 1, 2, 3 inputs, MBF(k), and several monotone Boolean models comprised of such MBFs.

We start by showing that there is a change of variables that results in a one-to-one correspondence between monotone Boolean models for any NL-free network and monotone Boolean models compatible with networks with all activating edges. We then analyze such positive networks and use the change of variables to obtain results for the original NL-free networks.

After illustrating our approach on a toggle switch network, we consider networks with a 2-team network structure, which were recently described [21, 22]. Here, edges connecting the nodes from the same team are activating, but those connecting nodes from opposite teams are repressing. We show that such networks are NL-free and that the most prevalent phenotypes are those where one team is active, while the other is not, and the most prevalent bistability is between these two phenotypes.

We then study two EMT networks: one a six-node network [2] and one with 15 nodes and 59 edges [7]. We show that the most prevalent states supported by both networks are the epithelial and mesenchymal states, and that bistability between them is the most prevalent bistability. This work also allows studying the prevalence of other steady states apart from mesenchymal and epithelial states, which are linked with the discussion about so-called *intermediate steady states*, which have biological significance [23].

The paper is organized as follows: After introducing basic definitions, we discuss negative-loop free (NL-free) networks in section *NL-free networks*. The main result of this section is that all monotone Boolean models of an NL-free network are in one-to-one correspondence with monotone Boolean models of a positive network. In section *Steady states for positive MBMs* we characterize MBM that support a particular steady state, and in sections *Bistability* and *Multistability*, those that support bistability and multistability, respectively. We illustrate our results throughout the text on several networks, including toggle switch, toggle switch with self-loops, 2-team network, and two EMT network models with 6 and 15 nodes. In the *Methods* section we use lattice theory to describe the sets of MBF, which we use to construct MBMs that support steady states. We close the paper with a discussion.

## 2 Results

### Definition 2.1

*A regulatory network RN* = (*V, E*, δ) *is a directed graph with nodes V with* |*V*| = *N, directed edges E, and an edge sign function* δ : *E* → {−1, 1}. *We denote an edge from node v*_*i*_ *to node v*_*j*_ *without indicating its sign by* (*v*_*i*_, *v*_*j*_) *or v*_*i*_⊸*v*_*j*_. *The edge v*_*i*_⊸*v*_*j*_ *is activating if* δ(*v*_*i*_, *v*_*j*_) = 1 *and repressing if* δ(*v*_*i*_, *v*_*j*_) = −1. *Graphically, an activating edge of a regulatory network edge is denoted by v*_*i*_ → *v*_*j*_ *and a repressing edge by v*_*i*_ ⊣ *v*_*j*_. *The sources and targets of a node v*_*i*_ *are given by*

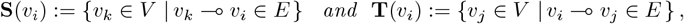

*respectively*.

### Definition 2.2

*A loop in network RN is a set of edges* (*v*_1_, *v*_2_), (*v*_2_, *v*_3_), …, (*v*_*k*_, *v*_*k*+1_) *with v*_1_ = *v*_*k*+1_. *A loop is positive (negative) if the product of the signs of edges*

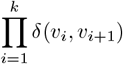

*is* 1 (−1).

### Definition 2.3

*A regulatory network RN* = (*V, E*, δ) *is negative loop free (NL-free), if it has no negative loops*.

### Definition 2.4

*A Boolean function f* : 𝔹^*k*^ → 𝔹 *is* increasing with respect to input *j if for any input b* = (*b*_1_, …, *b*_*k*_) ∈ 𝔹^*k*^

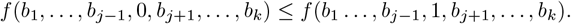

*It is* strictly increasing with respect to *j if there is at least one input b* ∈ 𝔹^*N*^ *where the inequality is strict*.

*A Boolean function f* : 𝔹^*k*^ → 𝔹 *is* decreasing with respect to input *j if for any collection of b*_*i*_ ∈ 𝔹

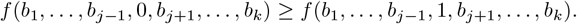

*It is* strictly decreasing with respect to *j if there is at least one input b* ∈ 𝔹^*k*^ *where the inequality is strict*.

### Definition 2.5

*A Boolean function f* : 𝔹^*k*^→ 𝔹 *is* monotone *if it is increasing or decreasing with respect to each input i* = 1, …, *k. In this case, f is called a* monotone Boolean function.

### Definition 2.6

*A set of increasing monotone Boolean functions f* : 𝔹^*k*^ → 𝔹 *will be denoted MBF* (*k*).

In Figure 1a we show three increasing monotone Boolean functions *MBF* (1) with single input denoted *X*. These are the zero function **0**, the identity function *Id*, and the one function **1**. We represent these three functions as nodes, connected by edges if their truth values differ on a single input string. In Figure 1b we list six increasing monotone Boolean functions *MBF* (2) with two inputs denoted *X* and *Y*. Their names are in the top row of the table; function *X* copies the values of the input *X*, while the function *Y* copies the input *Y*. Similarly, we represent these functions in the form of a graph where again, nodes are connected by edges if their truth values differ on a single input string.

**Fig 1.**
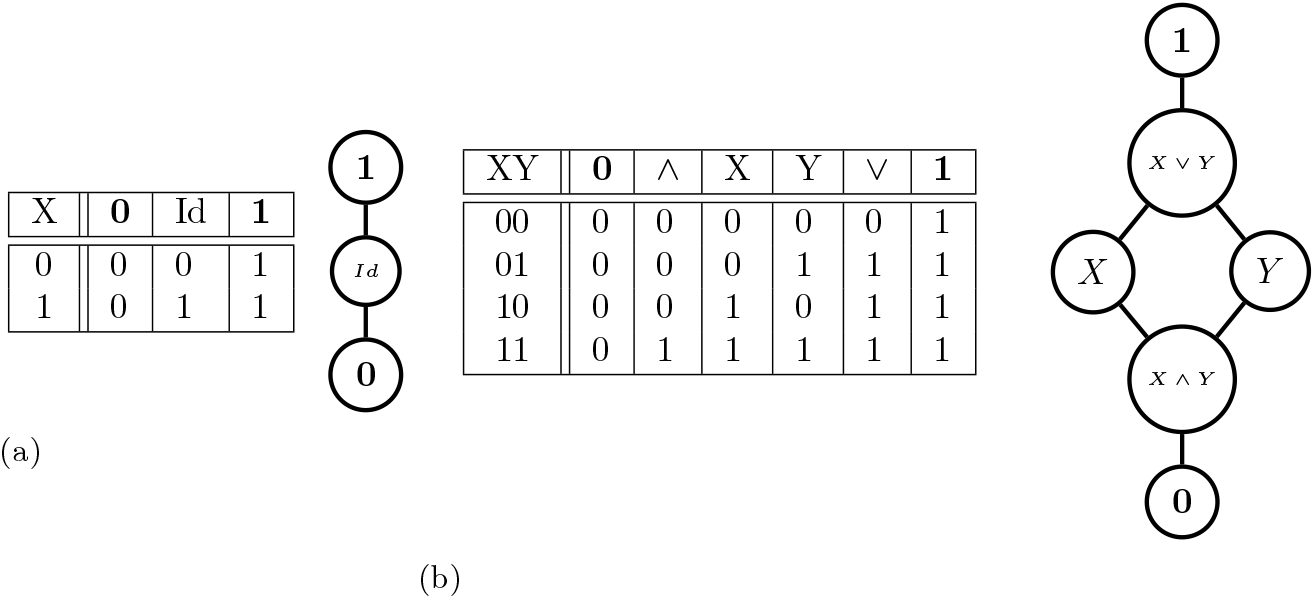
(a) There are three monotone Boolean functions with single input *MBF* (1) and (b) six monotone Boolean functions with 2 inputs *MBF* (2)

As we will see in section 3.1, the sets *MBF* (*k*) for any *k* form a mathematical structure called *lattice*. The language and results from lattice theory will help in understanding which Boolean models support which steady states.

We now discuss monotone Boolean models of network dynamics *f* = (*f*_1_, …, *f*_*N*_) where the update function *f*_*i*_ at the node *v*_*i*_ is a monotone Boolean function.

### Definition 2.7

*A Boolean model f* : 𝔹^*N*^ → 𝔹^*N*^, *f* = (*f*_1_, …, *f*_*N*_), *is* monotone *if for every i the function f*_*i*_ *is monotone*.

### 2.1 NL-free networks

Consider a monotone Boolean model *f* = (*f*_1_, …, *f*_*N*_). where each 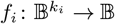. Apply a change of variables *α* = (*α*_1_, …, *α*_*N*_) where for each *i* either *α*_*i*_(*x*_*i*_) = *x*_*i*_ or *α*_*i*_(*x*_*i*_) = ¬*x*_*i*_. Let *E* ⊂ {1, …, *N* } be the indices where *α*_*i*_(*x*_*i*_) = *x*_*i*_.

Then

*i* ∈ *E* =⇒ *f*_*i*_ is increasing (decreasing) in *x*_*j*_ ↔ *f*_*i*_ is increasing (decreasing) in *α*(*x*_*j*_)

*i* ∈*/E* =⇒ *f*_*i*_ is increasing (decreasing) in *x*_*j*_ ↔ *f*_*i*_ is decreasing (increasing) in *α*(*x*_*j*_)

Observe that if *x*_*i*_ = *f*_*i*_(*x*) then ¬*x*_*i*_ = ¬*f*_*i*_(*x*) and *f*_*i*_ is increasing (decreasing) in *x*_*j*_ ↔ *α*(*f*_*i*_) is decreasing (increasing) in *x*_*j*_

#### Theorem 2.8.

*Consider NL-free network RN and a monotone Boolean model f* = (*f*_1_, …, *f*_*N*_) *compatible with RN*. *Then there is a change of variables α* : 𝔹^*N*^ → 𝔹^*N*^ *with α* ◦ *α* = *Id, and a positive Boolean model g* = (*g*_1_, …, *g*_*N*_) *such that the following diagram commutes*

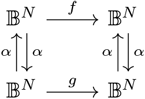

Proof can be found in section 3.1.

Note that the commutative diagram above implies that any trajectory under *f*, say

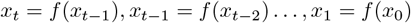

exists if, and only if there is a corresponding trajectory under *g*

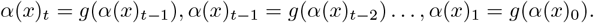

In particular, *x* is a fixed point under *f*, if, and only if *α*(*x*) is a fixed point under *g*

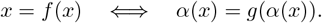

This shows that every NL-free network has the same dynamics as a positive network, where all edges are positive. Since an MBM *f* = (*f*_1_, …, *f*_*N*_) for a network with all positive edges has all functions *f*_*i*_ that are increasing MBFs, it is sufficient to study steady states and multistability for positive networks.

We illustrate the change of variables in Figure 2.

**Fig 2.**
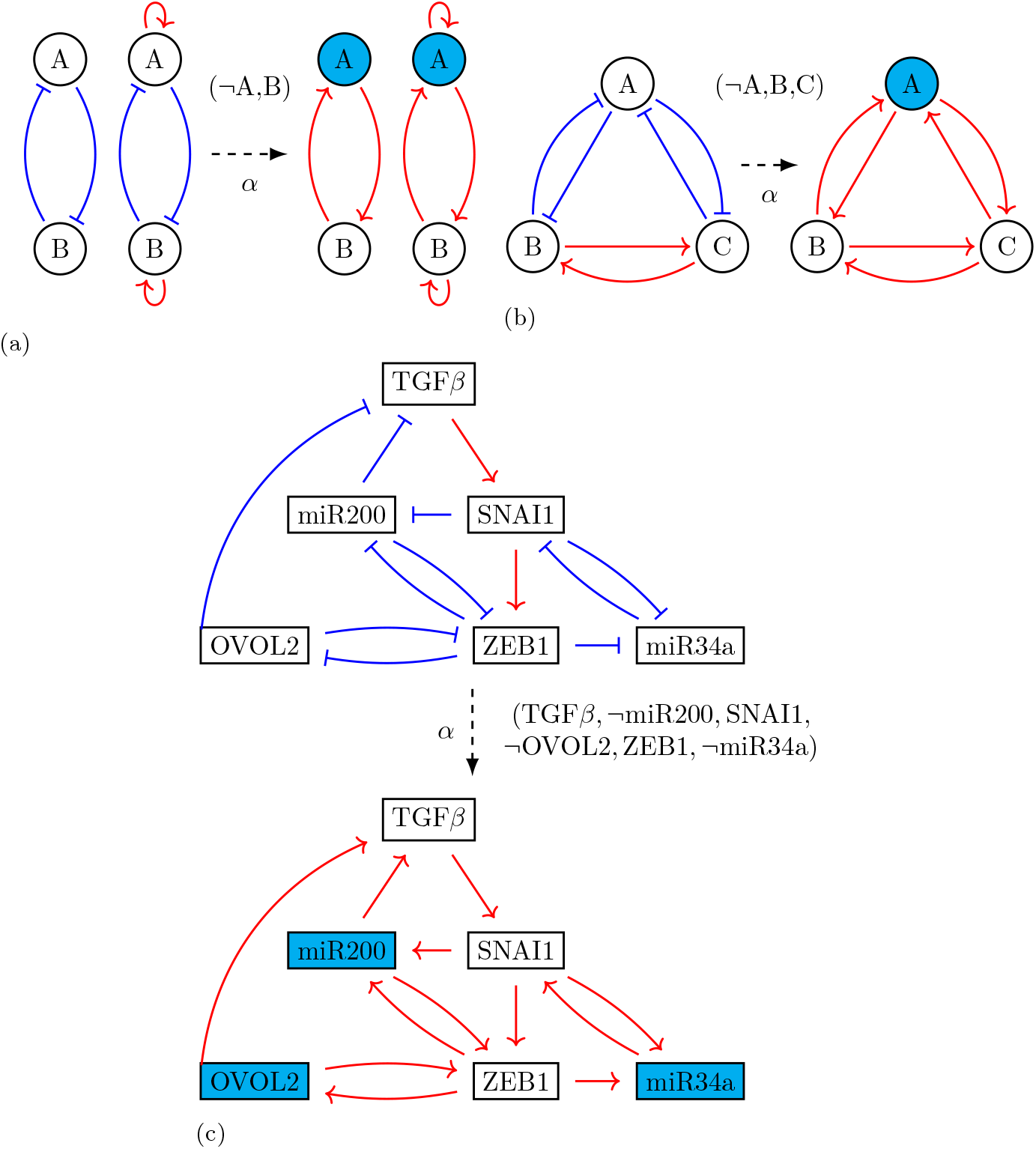
Schematic representation of the change of variables to obtain a positive network. The cyan nodes represent the nodes which have in *α*. (a): Toggle switch (left: without self-regulation, right: with self-activation), (b): 2-team network with 3 nodes, and (c): EMT network

In Figure 2a is a toggle switch [24], one of the basic network examples. Using Theorem 2.8, note that the map *α*(*A, B*) = (¬ *A, B*) results in both *g*_*B*_(*A*) and *g*_*A*_(*B*) to be increasing. Thus *α* = (¬ *A, B*) transforms *f* to a positive Boolean model *g* (Figure 2a). The change of variables *α* is not unique; it is easy to see that *α* = (*A*, ¬ *B*) also transforms *f* to a positive Boolean model *g*. Similarly, a toggle switch with self-activations can be transformed into a positive Boolean model using the same transformation.

In Figure 2b we consider a toy example of a 2-team network, studied in [25]. We define a 2-team network as a network where the set of nodes *V* can be decomposed into two disjoint sets *A* and *B, V* = *A*∪ *B*, in such a way that the nodes of the same team activate each other while inhibiting the nodes of the opposite team. In other words, *v*_*i*_ → *v*_*j*_ if, and only if, *v*_*i*_, *v*_*j*_ ∈ *A*, or *v*_*i*_, *v*_*j*_ ∈ *B*, and *v*_*i*_ *v*_*j*_, if, and only if *v*_*i*_ ∈ *A, v*_*j*_ ∈ *B*, or *v*_*i*_ ∈ *B, v*_*j*_ ∈ *A*.

Here node *A* belongs to one team, whereas *B* and *C* belong to the other team. It is easy to see that transformation *α* = (¬ *A, B, C*), or transformation *α* = (*A*, ¬ *B*, ¬ *C*) transforms *f* into a positive Boolean model *g* (Figure 2b).

Finally, we consider the biologically important *EMT* network which governs epithelial-to-mesenchymal transition (EMT) [2, 26], where we do not consider the input and output nodes from the original network and exclude the self-inhibition present on SNAI1. Then *α* = (TGF*β*, ¬ miR200, SNAI1, ¬ OVOL2, ZEB1, ¬ miR34a) transforms *f* into a positive model *g* (Figure 2c).

### 2.2 Steady states for positive MBMs

We are now ready to address the main question posed in this paper: *How many MBMs are compatible with an NL-free network support a particular steady state?*

Because of Theorem 2.8, it is sufficient to address this question only for positive monotone Boolean models. To this end, we define the following sets

#### Definition 2.9.

*Consider the set MBF(k), a set of monotone Boolean functions f* : 𝔹^*k*^ → 𝔹. *For any input b* ∈ 𝔹^*k*^ *let*

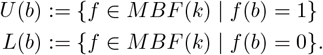

In other words, *U* (*b*)(*L*(*b*)) is the set of functions that evaluate to 1 (0). As an illustration, we show these intervals for the set MBF(2) of positive monotone Boolean functions (see Figure 1b) and Table 1.

**Table 1.** Sets of functions in MBF(2) that evaluate to 0 (left) and 1 (right). The third column in each part lists the number of functions in the corresponding set.

|  |  |  |  |  |  |
| --- | --- | --- | --- | --- | --- |
| L(11) | $\{\mathbf{0}\}$ | 1 | U(11) | $\{\mathbf{1}, X \vee Y, X, Y, X \wedge Y\}$ | 5 |
| L(10) | $\{\mathbf{0}, X \wedge Y, Y\}$ | 3 | U(10) | $\{\mathbf{1}, X \vee Y, X\}$ | 3 |
| L(01) | $\{\mathbf{0}, X \wedge Y, X\}$ | 3 | U(01) | $\{\mathbf{1}, X \vee Y, Y\}$ | 3 |
| L(00) | $\{\mathbf{0}, X \wedge Y, X, Y, X \vee Y\}$ | 5 | U(00) | $\{\mathbf{1}\}$ | 1 |

We are ready for the main result.

#### Theorem 2.10.

*Consider a positive network RN and an arbitrary Boolean state e* = (*e*_1_, …, *e*_*N*_) ∈ 𝔹^*N*^ *of the network RN*.

*Then e is a steady state of any monotone Boolean model f* = (*f*_1_, *f*_2_, …, *f*_*N*_) *where*

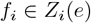

*Where*

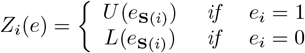

*where e*_**S**(*i*)_ *are the values of the input nodes of v*_*i*_ *evaluated at the state e*.

*Proof*. Vector *e* ∈ 𝔹^*N*^ is an equilibrium under a monotone Boolean model *f* = (*f*_1_, …, *f*_*N*_) where *f*_*i*_ ∈ *MBF* (|**S**(*i*)|) if and only if

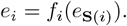

That is, *f*_*i*_ evaluated on the values of *e* restricted to sources of node *i* evaluates to the value *e*_*i*_. Since *U* (*e*_**S**(*i*)_) ⊂ *MBF* (|**S**(*i*)|) is the set of functions that evaluate to 1 on input *e*_**S**(*i*)_ and *L*(*e*_**S**(*i*)_) ⊂ *MBF* (|**S**(*i*)|) is the set of functions that evaluate to 0 on input *e*_**S**(*i*)_, the result follows.

□

In the section 3.1 we show that the sets *U* are ideals (down-sets) and filters (up-sets) in lattices. In addition, we show there the following characterization of the largest set *U* and the largest set *L*.

#### Lemma 2.11.

• *The largest set U* ⊂ *MBF* (*k*) *is the set U* (1_*k*_) *where* 1_*k*_ *is the vector of ones of length k, with* |*U* (1_*k*_)| = |*MBF* (*k*)| − 1

- *the largest set L* ⊂ *MBF* (*k*) *is the set L*(0_*k*_) *where* 0_*k*_ *is the vector of zeros of length k, with* |*L*(0_*k*_)| = |*MBF* (*k*)| − 1.

Proof can be found in section 3.1.

The next corollary of Lemma 2.11 characterizes the equilibria that are supported by most monotone Boolean models.

#### Theorem 2.12.

*Consider a positive network RN with N nodes. Then*

- *Every state in* 𝔹^*N*^ *is an equilibrium for some monotone Boolean model*.
- *For positive network RN with N among all the states, the states* 0 ∈ 𝔹^*N*^ *and* 1 ∈?𝔹^*N*^ *are most prevalent i*.*e. supported by most MBMs. The set of all MB models compatible with RN is*

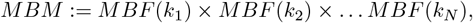

*where k*_*j*_ = |**S**(*v*_*j*_)|.

- *The sets of MBMs that supports* 0 ∈ 𝔹^*N*^ *has the form*

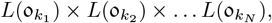

*which has the size*

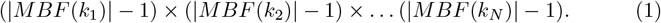

- *The sets of MBMs that supports* 1 ∈ 𝔹^*N*^ *has the form*

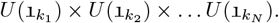

*which also has the size (1)*.

Proof can be found in section 3.1.

We are now ready to apply the theory to the examples in Figure 2. In the toggle switch, the change of variables *α* allows us to look at the equivalent problem of equilibria in a positive toggle switch where both edges are positive. Since each node has a single input, the set of MBMs *f* has the form *f* = (*f*_1_, *f*_2_) with *f*_*i*_ ∈ *MBF* (1). Therefore there are 3 *×* 3 = 9 monotone Boolean models *f* = (*f*_1_, *f*_2_), see Figure 1a for MBF(1) and Figure 3a for the set of models. It is easy to see that

**Fig 3.**
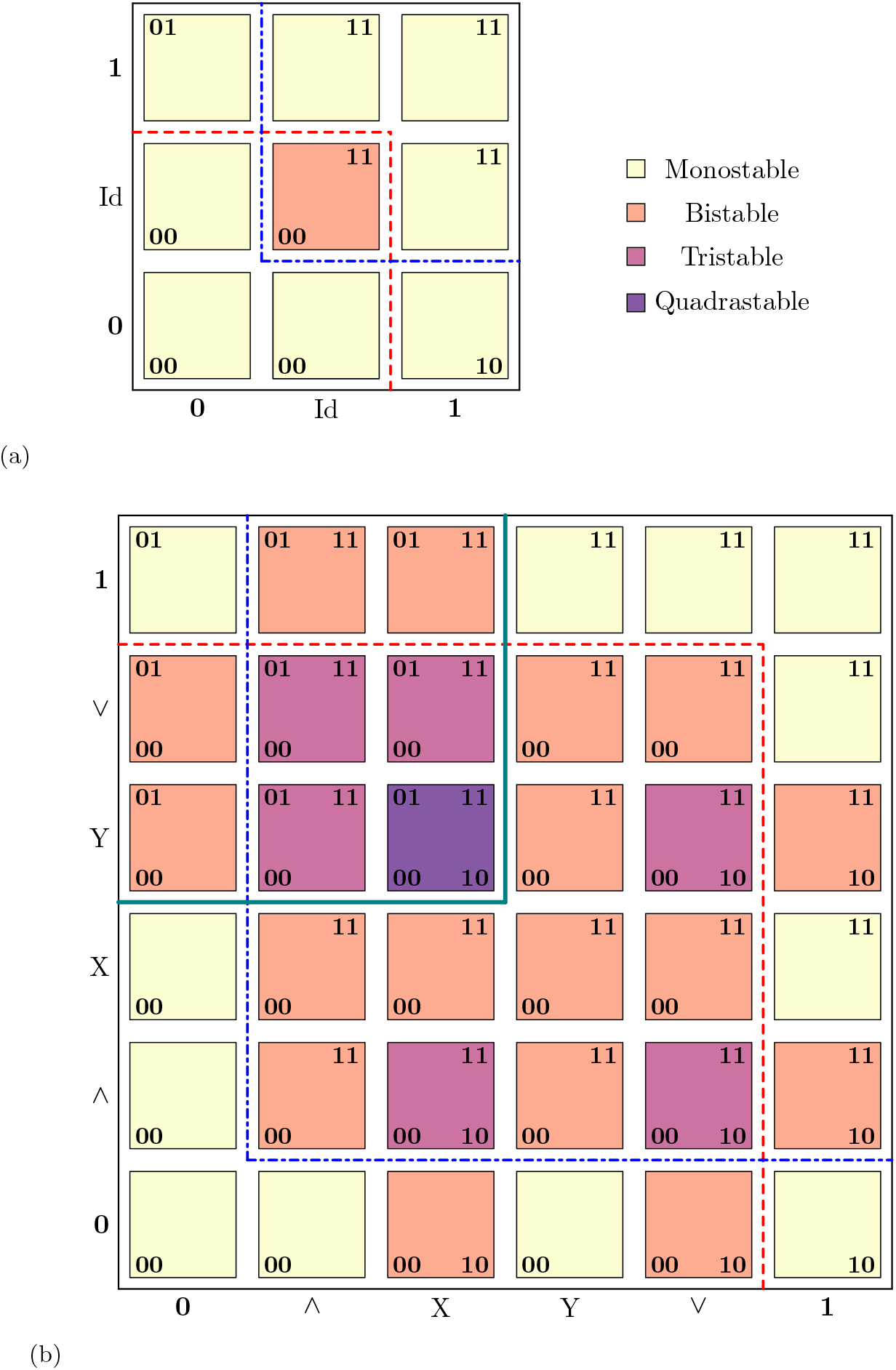
Parameter graphs (*f*_1_, *f*_2_) and the steady states supported by (a): *MBF* (1) *× MBF* (1) and (b): *MBF* (2) *× MBF* (2). The states supported are denoted in each square. The red dashed line, blue dash-dotted line, and the teal bold line delineate the set of models that support the steady states (00), (11), and (01), respectively. The filled colors represent the type of exact multistability supported by the corresponding Boolean model *f* = (*f*_1_, *f*_2_).

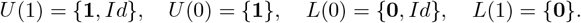

Therefore

- steady state (00) is supported by *f* ∈ *L*(0) *× L*(0) which has 4 models;
- steady state (10) is supported by *f* ∈ *U* (0) *× L*(1) which has 1 model;
- steady state (01) is supported by *f* ∈ *L*(1) *× U* (0) which has 1 model;
- steady state (11) is supported by *f* ∈ *U* (1) *× U* (1) which has 4 models.

Applying the change of variable function *α* : 𝔹^2^→ 𝔹^2^ of the form (*A, B*) → (¬ *A, B*) to the equilibria set in this list, we have the following result for the toggle switch

- steady state (10) is supported by 4 models;
- steady state (00) is supported by 1 model;
- steady state (11) is supported by 1 model;
- steady state (01) is supported by 4 models.

For the toggle switch with self-activation, the same change of variables *α* as in the toggle switch changes the model to *g* = (*g*_1_, *g*_2_) where each *g*_*i*_ ∈ *MBF* (2) has two inputs, see Figure 1b The set of MBMs has 6 *×* 6 = 36 models, see Figure 3b for the set of models and Table 1 for the list of intervals *U* (*b*), *L*(*b*) ⊂ *MBF* (2). The dominant steady states are the same as in the toggle switch network, but they are supported by a larger number of models:

- steady state (00) is supported by *f* ∈ *L*(00) *× L*(00) which has 5 *×* 5 = 25 models;
- steady state (11) is supported by *f* ∈ *U* (11) *× U* (11) which also has 5 *×* 5 = 25 models.

After applying *α*, the most prevalent steady states in the toggle switch with self-activation are

- steady state (10) supported by 25 models;
- steady state (01) supported by 25 models.

For the 2-team network in Figure 2b, any MBM *f* = (*f*_1_, *f*_2_, *f*_3_) can be changed to *g* = (*g*_1_, *g*_2_, *g*_3_) with *g*_*i*_∈ *MBF* (2). Again, all states are supported by some models, but the most prevalent steady states are

- steady state (000) is supported by *f* ∈ *L*(00) *× L*(00) *× L*(00) which has 5^3^ = 125 models;
- steady state (111) is supported by *f ∈ U* (11) *× U* (11) *× U* (11) which also has 5^3^ = 125 models.

Applying *α* : 𝔹^3^ → 𝔹^3^, *α* = (*A, B, C*) → (¬ *A, B, C*), gives the most common steady states as

- (100) supported by 125 models;
- (011) supported by 125 models.

Finally, for the EMT network in Figure 2c the change of variables

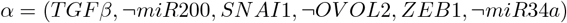

transforms *f* into a positive model *g* (Figure 2c). The collection of MBMs *f* = (*f*_*β*_, *f*_200_, *f*_*S*_, *f*_*O*_, *f*_*Z*_, *f*_34_) for the positive EMT (P-EMT) network is

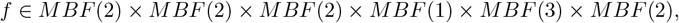

where the numbers correspond to the number of inputs of the corresponding nodes. The collection *MBF* (3) is shown in Figure 6 and has 20 functions. Therefore, there is a total of

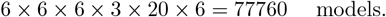

Within this collection of MBMs, the most common steady states are

- (111111) which is supported by models in 5 *×* 5 *×* 5 *×* 2 *×* 19 *×* 5 = 23750 models;
- which is also supported by 23750 models.

Applying the change of variables *α*, it follows that the most common steady states in the original EMT network are states

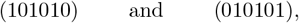

each of which is supported by 23750 out of 77760 MBMs compatible with the EMT network. These two states represent the mesenchymal and epithelial state, respectively, with the mesenchymal state characterized by TGF*β*, SNAI1, ZEB1 high, whereas the epithelial state is characterized by miR200, OVOL2, miR34a high.

Similarly, Theorem 2.10 can be used to calculate the number of MBMs supporting the other states for the positive network, as given in Table 2. These states correspond to the intermediate states characterized by mixed expression of the canonical epithelial and mesenchymal markers.

**Table 2.** Number of models supporting the top 12 most common steady states. P-EMT-state represents the state in the positive EMT network and EMT state represents the state in the original network after transformation.

| P-EMT state | Parameter sets | No. of MBMs | EMT state |
| --- | --- | --- | --- |
| (000000) | $L(00) \times L(00) \times L(00) \times L(0) \times L(000) \times L(00)$ | 23750 | (010101) |
| (111111) | $U(11) \times U(11) \times U(11) \times U(1) \times U(111) \times U(11)$ | 23750 | (101010) |
| (000100) | $L(01) \times L(00) \times L(00) \times U(0) \times L(001) \times L(00)$ | 5250 | (010001) |
| (111011) | $U(10) \times U(11) \times U(11) \times L(1) \times U(110) \times U(11)$ | 5250 | (101110) |
| (001001) | $L(00) \times L(10) \times U(01) \times L(0) \times L(010) \times U(10)$ | 3780 | (011100) |
| (110110) | $U(11) \times U(01) \times L(10) \times U(1) \times U(101) \times L(01)$ | 3780 | (100011) |
| (100100) | $U(01) \times L(00) \times L(10) \times U(0) \times L(001) \times L(00)$ | 3150 | (110001) |
| (011011) | $L(10) \times U(11) \times U(01) \times L(1) \times U(110) \times U(11)$ | 3150 | (001110) |
| (000001) | $L(00) \times L(00) \times L(01) \times L(0) \times L(000) \times U(00)$ | 2850 | (010100) |
| (111110) | $U(11) \times U(11) \times U(10) \times U(1) \times U(111) \times L(11)$ | 2850 | (101011) |
| (100000) | $U(00) \times L(00) \times L(10) \times L(0) \times L(000) \times L(00)$ | 2850 | (110101) |
| (011111) | $L(11) \times U(11) \times U(01) \times U(1) \times U(111) \times U(11)$ | 2850 | (001010) |

Consider now a larger EMT network in Figure 4, described in [7], with 15 nodes and 59 edges (excluding the input/output nodes and self-inhibitions). The description of sets *L*(·), *U* (·) becomes computationally challenging due to nodes like ZEB1 and ZEB2 having many inputs (12 and 10, respectively). However, leveraging Theorem 2.12 allows identification of the most common states by transforming the most common states 0 and 1 of the positive network using *α*. It can be seen that this network is NL-free, and there exists *α* given by:

**Fig 4.**
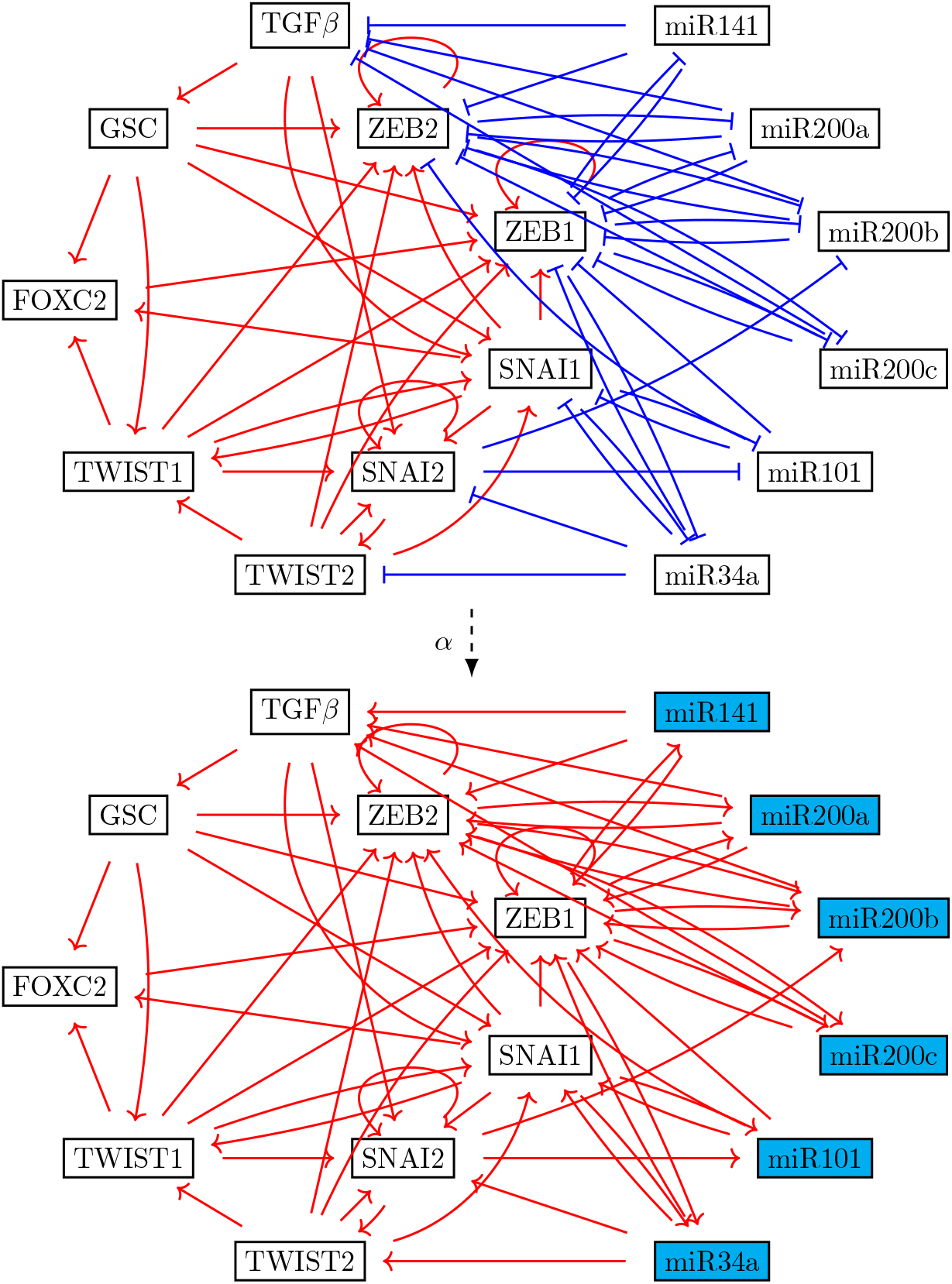
Schematic representation of the change of variables to obtain a positive network of EMT network from [7]. The cyan nodes represent the nodes which have ¬ in *α*.

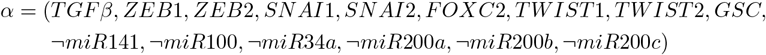

Therefore the most common states in the network in Figure 4 are (1) TGF*β*, ZEB1, ZEB2, SNAI1, SNAI2, FOXC2, TWIST1, TWIST2, GSC high and miR141, miR100, miR34a, miR200a, miR200b, miR200c low, which corresponds to the mesenchymal state and (2) the state where the first group of nodes are low and the second group is high, which corresponds to the epithelial state. This mathematical result supports the biological insight that the main role of this model is to modulate the transition between these two states. At the same time, the fact that the most prevalent states in the MB models are indeed the epithelial and mesenchymal states supports the idea that looking for the most prevalent states in MB models of a network may reveal the biological significance of the network.

### 2.3 Bistability

As we have shown in Theorem 2.12, the most common equilibria in any NL-free network have the same abundance. A natural question arises whether the bistability between these two equilibria is also the most common form of bistability. Note that in our work,c we will refer to the presence of two or more steady states by the term *bistability*. We will use the term *exact bistability* for the presence of exactly two steady states. Specifically, when we say that a model supports bistability between states *d* and *e*, we mean that *d* and *e* are among the steady states, but other steady states beyond *d* and *e* may also exist.

The following theorem extends Theorem 2.10 by characterizing the MBMs that support a bistability between two steady states in the set of positive MBMs.

#### Theorem 2.13.

*Consider a positive network RN and two arbitrary Boolean states d, e* ∈ 𝔹^*N*^. *Then the bistability between steady states d and e is supported by any positive monotone Boolean model f* = (*f*_1_, *f*_2_, …, *f*_*N*_) *where*

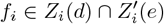

*where*

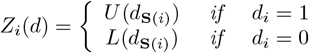

*and*

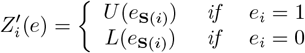

*where d*_**S**(*i*)_, (*e*_**S**(*i*)_) *are the values of the input nodes of v*_*i*_ *evaluated at the state d*, (*e*), *respectively*.

There are several important consequences of this description. If *i*-th components of *d* and *e* agree, *d*_*i*_ = *e*_*i*_ then

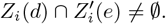

However, if *d*_*i*_ ≠ *e*_*i*_ then the intersection may be empty. Since *d* and *e* must differ in at least one component, not all pairs of states *d, e* can be bistable states for an MBM.

For instance, in the positive toggle switch the bistability between *d* = (00) and *e* = (11) is supported by the set of MBMs *f* = (*f*_1_, *f*_2_) with

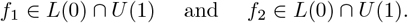

These intersections are non-empty and contain a single model

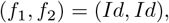

see Figure 3a. This corresponds to a single MBM supporting bistability between (01) and (10) in the toggle switch. However, bistability between (00) and (10) in *MBF* (1) *× MBF* (1) is supported by models *f* = (*f*_1_, *f*_2_) with

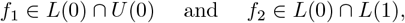

where *L*(0) ∩ *U* (0) = ∅. Applying the map *α*, we conclude there are no MBMs for the toggle switch supporting bistability between (10) and (00). In fact, no other pair of states except (01) and (10) can be bistable in a toggle switch, see Figure 3a.

In the case of a toggle switch with self-activation, a larger set of models offers bistability between the states (00) and (11) in the positive network, see Figure 1b. These are the set of MBMs

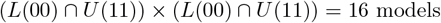

Hence, bistability between (01) and (10) is supported by 16 models for the toggle switch with self-activation. However, unlike the toggle switch, bistability between the other pairs of steady states is also supported. The number of models that support these bistabilities in the positive toggle switch are

- and (01): 6 models;
- and (10): 6 models;
- and (11): 6 models;
- (10) and (11): 6 models;
- and (10): 1 model.

#### Lemma 2.14.

*Consider a positive network RN with N vertices. Then the most common bistability supported by MBMs f* = (*f*_1_, …, *f*_*N*_) *with*

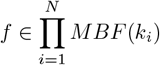

*is between the steady states d* = 0 *and e* = 1.

*It is supported by*

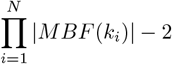

*MBMs*.

Proof can be found in section 3.1.

For 2-team network example, Lemma 2.14 shows that the bistability between (100) and (011) is supported by 4^3^ = 64 MBMs, which is 64*/*216 = 0.296; or almost 30% of all models.

For the 6-node EMT network, the bistability between epithelial and mesenchymal states is supported by 4 *×*4 *×* 4 *×* 1 *×* 18 *×* 4 = 4608 MBMs. The resulting prevalence is 4608*/*77760 = 0.059, or about 6%.

### 2.4 Multistability

Mirroring our comment about bistability, we will say that an MB model supports *q* − multistability if it has *n* ≥ *q* steady states. We will use the term *exact q-multistablity* for MBM that supports exactly *q*-steady states.

In order to determine the collection of MBMs that support exact *q*−multistability, it is sufficient to know MBMs that support *q*−multistability for all *n* ≥ *q*. More precisely, let *m*_*n*_(*E*) be the set of MB models that support *n*−multistability between states *E* := {*e*_1_, …, *e*_*n*_}, for some *n* ≥ *q*, and let *em*_*q*_(*A*) be the set of MBMs that support exact *q*−multistability between states *A* = {*e*_1_, …, *e*_*q*_}. Then

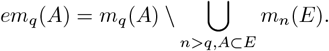

The extension of Theorems 2.10 and 2.13 to multistability is straightforward.

#### Theorem 2.15.

*Consider a positive network RN and q Boolean states d*^1^, …, *d*^*q*^ ∈ 𝔹^*N*^. *Then the multistability between these steady states is supported by any positive monotone Boolean model f* = (*f*_1_, *f*_2_, …, *f*_*N*_) *where*

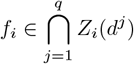

*where*

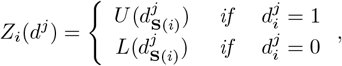

*where* 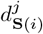 *are the values of the input nodes of v*_*i*_ *evaluated at the state d*^*j*^, *respectively*.

The conditions under which these products are non-empty is subject to ongoing investigation.

As an example, note that in Figure 3b there is one set of functions (*f*_1_, *f*_2_) that supports multi-stability between all 4 states (00, 10, 01, 11), 4 models that support tristability between (00, 01, 11) and 4 models that support tristability between (00, 10, 11).

The question of what is the highest multistability supported by the network is related to the number of positive loops in the network, and only partial solutions and estimates are known [27].

## 3 Methods

### 3.1 Lattice theory

#### Definition 3.1

([28]). *A partially ordered set P (or poset) is a set P together with a binary relation denoted* ≤, *satisfying the following three axioms:*

*1. For all t* ∈ *P, t* ≤ *t (reflexivity)*.

*2. If s* ≤ *t and t* ≤ *s, then s* = *t (antisymmetry)*.

*3. If s* ≤ *t and t* ≤ *u, then s* ≤ *u (transitivity)*.

*We say that two elements s and t of P are comparable if s* ≤ *t or t* ≤ *s; otherwise s and t are incomparable, denoted s* ║*t*.

#### Definition 3.2.

*A subset D of a poset P is a down-set or order ideal if*

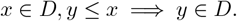

*We denote the down-sets in P by J* (*P*). *Dually, a subset U of P is an up-set or order filter if*

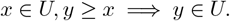

*We denote the up-sets of P by U* (*P*). *Since we will only deal with order ideals and order filters, we will shorten these terms to ideal and filter, respectively*.

#### Definition 3.3.

*Let P be a poset with two binary operations* ∨, *called join, and*∧, *called meet. P is a lattice if every pair of elements of P has both a meet and a join*.

In this paper, we will only consider finite posets and lattices. In a finite lattice, all ideals (filters) are principal [29] and thus have the form

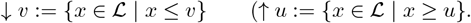

#### Definition 3.4.

*Let* ℒ *be a lattice*.

*1. A nonzero element j is join-irreducible if j is not the join of two smaller elements, that is, if*

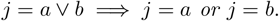

*2. Dually, a nonunit element m* ∈ ℒ *is meet-irreducible if m is not the meet of two larger elements, that is, if*

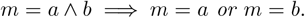

A subset *A* ⊊?*B* is a *proper* subset of *B*.

#### Definition 3.5

([29]). *Let* ℒ *be a lattice*.

*A proper ideal J is prime if*

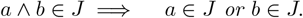

*Dually, a proper filter F is prime if*

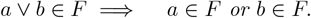

#### Theorem 3.6

(Theorem 3.37, [29]). *The properties of being a prime ideal and being a prime filter are complementary, that is, I is a prime ideal if and only if I*^*c*^ *is a prime filter*.

#### Definition 3.7

([29]). *If a lattice* ℳ *has a smallest element (which we call* 0*), and* ℒ *is another lattice then the ideal kernel of a lattice homomorphism f* : ℒ → ℳ *is the set*

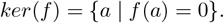

*This is easily seen to be an ideal of* ℒ.

Given a lattice ℒ, choosing ℳ= 𝔹 to be the Boolean lattice 𝔹 := 𝔹 (0, 1 ; *<*), we arrive at the concept of the indicator function of a subset *S* ⊆ ℳ. The *indicator function* of *S* is the function *f*_*S*_ : ℒ →𝔹 defined by setting *f*_*S*_(*x*) = 0 if and only if *x* ∈ *S*. In general, an indicator function need not be a lattice homomorphism. However, the indicator function of a **prime** ideal *P* is a lattice epimorphism.

#### Definition 3.8

([28]). *Lattice* ℒ *is a distributive lattice if*

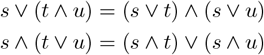

*holds for any elements s, t, u* ∈ ℒ.

The fundamental Theorem for finite distributive lattices is due to Birkhoff [30].

#### Theorem 3.9

(Birkhoff Theorem). *Any finite distributive lattice is isomorphic to the lattice of down-sets of its join-irreducible elements*.

This representation is crucial for understanding the relationship between join-irreducibles and prime ideals.

#### Lemma 3.10.

*Down set of join-irreducible element in finite distributive lattices are prime ideals*.

*Proof*. Consider an ideal (↓ *v*). If *x* ∨ *y* ∈ (↓ *v*), then by join-irreducibility of *v*, either *x* ≤ *v* or *y* ≤ *v*, and therefore that element is also in ↓ *v*. This directly satisfies the definition of a prime ideal.

#### Theorem 3.11

(Theorem 3.41, [29]). *In a lattice* ℒ, *the prime ideals in* ℒ *are precisely the kernels of the indicator epimorphisms*.

By duality Theorem 3.6, the prime filters are precisely sets *U* which map to 1 under an indicator epimorphism.

### 3.2 Lattice of increasing monotone Boolean functions

Consider the poset 𝔹^*k*^ of Boolean vectors under the order induced by 0 *<* 1 on each component.

The set of increasing monotone Boolean functions *MBF* (*k*), defined in Definition 2.6, is a set of order preserving maps

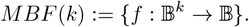

#### Definition 3.12

(Truth set and zero set). *The truth set T* (*f*) *of a Boolean function f* : 𝔹^*k*^ → *𝔹 is*

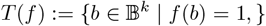

*while the zero set* ker(*f*) *is*

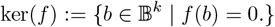

Note that (*MBF* (*k*), ⪯) is a poset with order given by

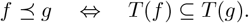

#### Lemma 3.13.

*The poset* (*MBF* (*k*), ⪯) *is a distributive lattice with operations given by union and intersection of the associated truth sets*

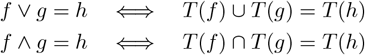

Let *G* : *MBF* (*k*) → *U* (*P*) taking *f* → *T* (*f*)

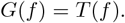

The map *G* is a lattice isomorphism between (*MBF* (*k*), ⪯) and (*U* (*P*), ⊆) where we made the orders explicit.

We also define an *anti-order* ⪯_*a*_ on *MBF* (*k*) by using kernels of the maps *f*

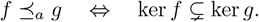

It is straightforward to see that

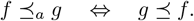

Consider a map *F* : (*MBF* (*k*), ⪯_*a*_) → *J* (*P*) taking *f* → ker *f* and so

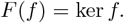

Here *J* (*P*) is the set of order ideals in *P*. It is easy to see that the map *F* is an isomorphism and thus (*MBF* (*k*), ⪯_*a*_) ≅ *J* (*P*).

We have an immediate result.

#### Lemma 3.14.

(*MBF* (*k*), ⪯_*a*_) *is a lattice with operations* (∨_*a*_, ∧_*a*_)

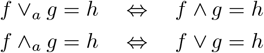

*where the operations* (∨, ∧) *are operations in lattice* (*MBF* (*k*), ⪯)

We will use notation *MBF* (*k*) for (*MBF* (*k*), ⪯) and notation *MBF*^*a*^ for lattice (*MBF* (*k*), ⪯ _*a*_).

Let *S* = 2^*P*^ be a set of all subsets of finite poset *P*. Let

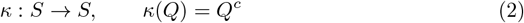

which assigns to *Q* ∈ *S* its complement *Q*^*c*^ in *S*. Then the map *κ* applied to poset *P* := 𝔹^*k*^ of all possible inputs to *f* ∈ *MBF* (*k*) maps *T* (*f*) to ker *f* and ker *f* to *T* (*f*).

#### Corollary 3.15.

*The map κ* : *P* → *P induces an isomorphism between lattices* (*MBF* (*k*), ⪯) *and* (*MBF*^*a*^(*k*), ⪯_*a*_) *which maps meet-irreducible elements to join-irreducible elements and join-irreducible elements to meet-irreducible elements. The isomorphism takes each f* ∈ *MBF* (*k*) *to itself but exchanges the lattice operations*.

*Proof*. Since for any two functions *f, g* ∈ *MBF* (*k*) the union *T* (*f*) ∪ *T* (*g*) is the complement of ker *f* ∩ ker *g* and intersection *T* (*f*) ∩ *T* (*g*) is the complement of ker *f* ∪ ker *g* by de Morgan laws, the result follows from Lemma 3.14.

Recall that in Definition 2.9 we set

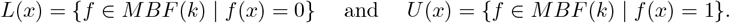

#### Theorem 3.16.

*We have the following characterizations of the sets L*(*x*), *U* (*x*); *x* ∈ 𝔹^*k*^ :

*1. For each x the set L*(*x*) *is a prime ideal in MBF* (*k*).

*2. There is a unique meet-irreducible element g* = *g*(*x*) ∈ *MBF* (*k*) *such that L*(*x*) *is a down-set of g in MBF* (*k*), *i*.*e. L*(*x*) =↓ *g*.

*3. For each x the set U* (*x*) *is a prime filter in MBF* (*k*)

*4. There is a unique join-irreducible element g* = *g*(*x*) ∈ *MBF* (*k*) *such that U* (*x*) *is an up-set of g in MBF* (*k*), *i*.*e. U* (*x*) =↑ *g*.

*Proof*. For illustration of the arguments below, please see Figure 5 for MBF(2). The set MBF(3) is shown in Figure 6.

**Fig 5.**
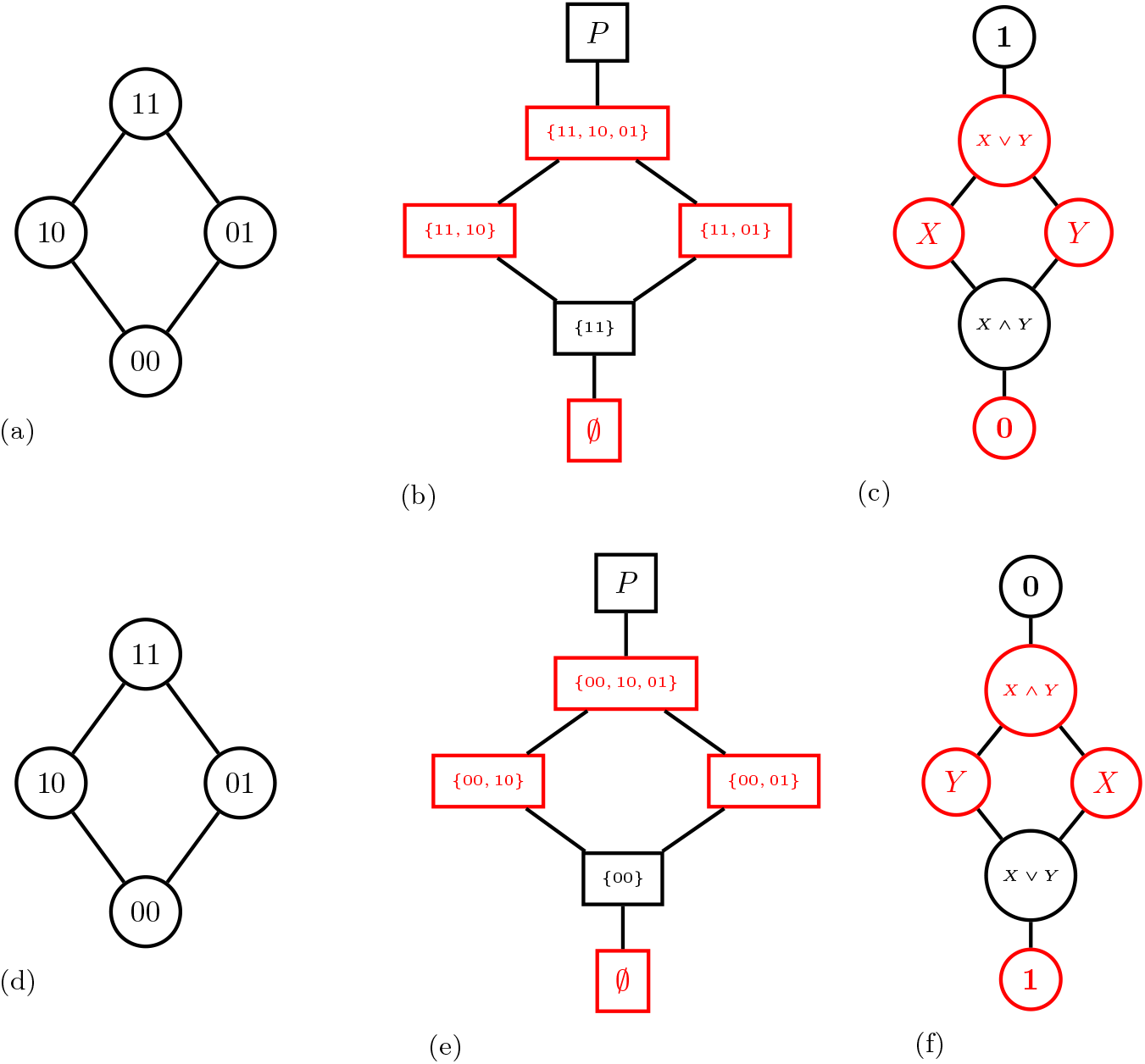
(a) Poset *P* ; (b) Lattice of up-sets of *U* (*P*); (c) The set of monotone Boolean functions with two inputs *MBF* (2) is isomorphic to *U* (*P*) via the map *f* → *T* (*f*). (d) Poset *P* ; (e) Lattice of down-sets of *J* (*P*); (f) The set of monotone Boolean functions with two inputs *MBF*^*a*^(2) with anti-order is isomorphic to *J* (*P*) via the map *f* → ker *f*.

**Fig 6.**
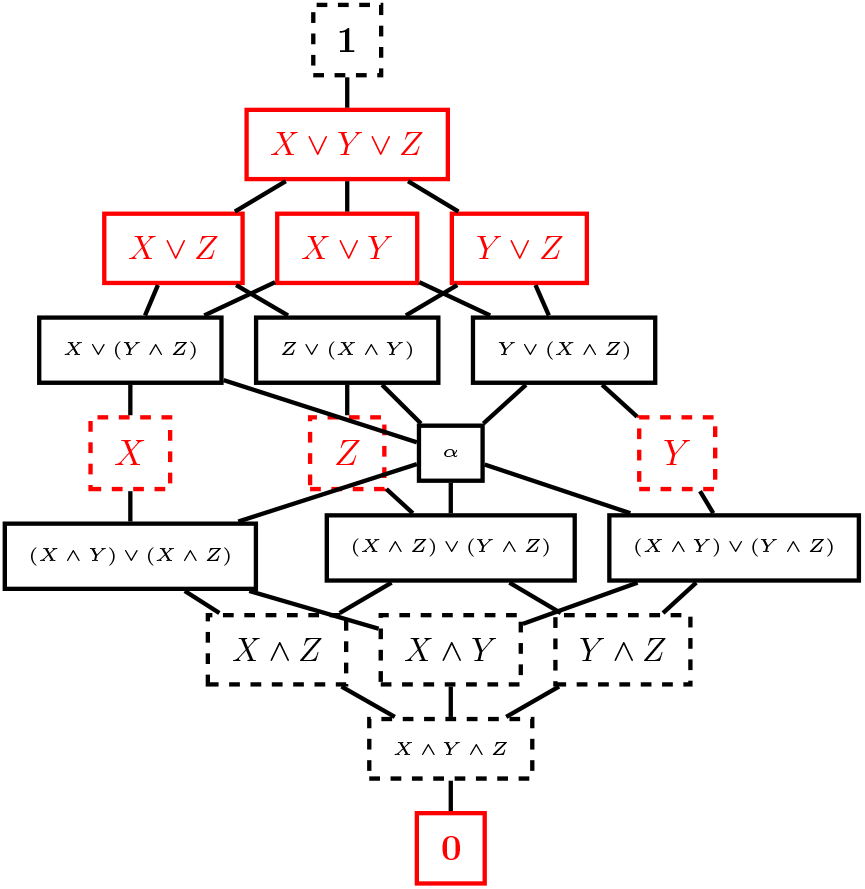
Lattice of monotone Boolean functions *MBF* (3). Nodes aligned horizontally have the same size of truth set, ranging from 0 at the bottom to 8 = 2^3^ on the top. Each node has DNF description of the function, where **0** and **1** are constant functions and *α* = (*X* ∧ *Z*) ∨ (*X* ∧*Y*) ∨ (*Y* ∧ *Z*). Down-sets of meet-irreducible nodes (red) are the sets *L*(∗); while up-sets of join-irreducible elements (dashed) are the sets *U* (∗). Both collections are isomorphic to the poset *P* = 𝔹^3^.

We start by describing all epimorphisms *U* (*P*) → 𝔹. First, for every *x* ∈ *P* let

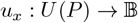

be a bounded lattice homomorphism which maps all up-sets of *P* containing *x* to 1 and all other up-sets to 0. Note that this set of epimorphisms is in one-to-one correspondence with *x* ∈ *P*.

We now show that every lattice epimorphism *f* : *U* (*P*) → 𝔹 has the form *u*_*x*_ for some *x* ∈ *P*. Consider any lattice epimorphism *f* : *U* (*P*) → 𝔹. Then the elements *y* ∈ *U* (*P*) such that *f* (*y*) = 0 that are mapped to 0 must have a unique maximal element 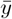, which is the join of the elements *y* that are mapped to 0. This follows from the homomorphism property requiring that if *f* (*u*) = 0 and *f* (*v*) = 0, then also *f* (*u* ∨ *v*) = 0. Importantly, 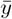 must be meet-irreducible, since it cannot be the meet *u v* of any elements *u*, ∧ *v* with *f* (*u*) = *f* (*v*) = 1.

Since *U* (*P*) ≅ (*MBF* (*k*), ⪯) and *J* (*P*) ≅ *MBF*^*a*^(*k*), ⪯_*a*_), the isomorphism from Corollary 3.15 between lattices

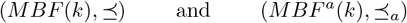

induces isomorphism between *U* (*P*) and *J* (*P*), By Lemma 3.14 the meet irreducible elements in *U* (*P*) are mapped to join-irreducible elements in *J* (*P*). Since the isomorphism between (*MBF* (*k*), ⪯) and (*MBF*^*a*^(*k*), ⪯_*a*_) is induced by *κ*, it follows from (2) that 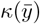 is a join irreducible element of *J* (*P*).

We illustrate this map on the example *MBF* (2) in Figure 5. The set *U* (*P*) is depicted in panel Figure 5b and the red nodes correspond to meet-irreducible elements of *U* (*P*). Note that *κ*({11, 10, 01}) = {00}, *κ*({11, 10}) = {00, 01 }, *κ*({ 11, 01}) = {00, 10} and *κ*({ ∅} = *P* and the images are join-irreducible nodes of the lattice *J* (*P*) in Figure 5e.

By Lemma 3.10 down-sets of join-irreducible elements in *J* (*P*) are prime ideals. Therefore for each 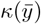 there exists 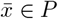 such that

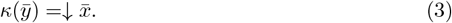

To summarize our construction so far, for each lattice epimorphism *f* : *U* (*P*) → 𝔹we associate a meet-irreducible element 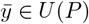, to which we assign a join-irreducible element 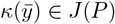, which has a form of (3) form some 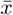. We claim that 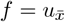. First, we note that the kernel of the epimorphism *f* are all the up-sets that are subsets of the largest up-set 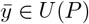. By construction, the set 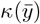 is the complement of the up-set 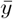 in *S* = 2^*P*^ and so it is disjoint from any up-sets in *U* (*P*) that are below 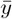. Furthermore, this complement 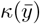 is a prime ideal generated by 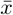. Therefore, 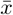 is the largest element in *P* which does not belong to the upper set 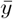. As a consequence, if *y* ∈ *U* (*P*) contains 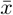, then *f* (*y*) = 1, but if 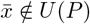, then *f* (*y*) = 0. This shows that 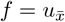.

Therefore, every lattice epimorphism *f* has the form *f* = *u*_*x*_ for some *x*. Consider *u*_*x*_ : *MBF* (*k*) → 𝔹 for some *x* ∈ 𝔹^*k*^. Then the set of up-sets in *P* that do not contain *x* is a down-set in *MBF* (*k*) that consists of all functions *f* ∈ *MBF* (*k*)

whose truth set does not contain *x*, and therefore *f* (*x*) = 0. By definition of the sets *L*(*x*), these are precisely the functions that belong to the set *L*(*x*). This down-set in *MBF* (*k*) is mapped by *u*_*x*_ to 0 and therefore is a kernel of a homomorphism *u*_*x*_. By Theorem 3.11 *L*(*x*) is the kernel of indicator epimorphism and therefore prime ideal in *MBF* (*k*). Furthermore there is unique maximal element 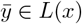 which is meet -irreducible and such that

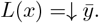

This proves statements 1 and 2 of the Theorem.

We illustrate this in Figure 5c. Down-sets of meet-irreducible elements (in red) are *L*(*X* ∨ *Y*) = *L*(00), *L*(*Y*) = *L*(01), *L*(*X*) = *L*(10), *L*(**0**) = *L*(11). At the same time *L*(11) = ker *u*_00_ = {**0**}, *L*(10) = ker *u*_01_ = {**0**, *X* ∧ *Y, X*}, *L*(01) = ker *u*_10_ = {**0**, *X* ∧ *Y, Y* } and *L*(00) = ker *u*_11_ = {**0**, *X* ∧ *Y, X, Y, X* ∨ *Y* }.

By Theorem 3.6 the complements of prime ideals are prime filters. This shows *U* (*x*) are upsets of join-irreducible elements which proves 3 and 4.

To illustrate, note that in Figure 5f the down-sets of meet-irreducible elements in *MBF*^*a*^(2) (in red), which are upsets of join-irreducibles in *MBF* (2), are *U* (*X* ∧ *Y*) = *U* (11), *U* (*Y*) = *U* (01), *U* (*X*) = *U* (10), *U* (**1**) = *U* (00).

### 3.3 Structure of MBF models that support fixed points

In this subsection, we use the lattice theory language to describe the sets *L*(*x*), *U* (*x*) ⊂ *MBF* (*k*).

#### Proof of Lemma 2.11

*Proof*. By Theorem 3.16, each set *U* (*x*) is a prime filter in MBF(k). Since the logical function in *k* arguments *X*_*i*_ given by

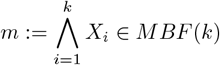

has unique predecessor **0** ∈ *MBF* (*k*), ↑ *m* is joint-irreducible element in *MBF* (*k*). Consequently, *m* is a prime filter and it contains all functions in MBF(k), except the function **0**. Therefore, this is the maximal filter with size |*MBF* (*k*) |−1. Finally, since **0** is the only function that evaluates to 0 on input *b* = − 1, it follows that ↑ *m* = *U* (1).

Analogously, by Theorem 3.16 each set *L*(*x*) is a prime ideal in MBF(k). The logical function in *k* arguments *X*_*i*_ given by

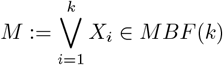

has unique successor **1** ∈ *MBF* (*k*), *M* is meet-irreducible element in *MBF* (*k*) and ↓ *M* is a prime ideal that contains all functions in MBF(k), except the function **1**. Therefore, this is the maximal ideal with size |*MBF* (*k*)| − 1. Finally, since **1** is the only function that evaluates to 1 on input 0, it follows that ↓ *M* = *L*(0).

#### Proof of Theorem 2.12

*Proof*. Any state *b* ∈ 𝔹^*N*^ is supported by the monotone Boolean model *f* = (*f*_1_, …, *f*_*N*_) where *f*_*i*_ is a constant function *f*_*i*_ = *b*_*i*_. Therefore, the set of models supporting *b* is non-empty.

The rest of the Theorem follows by realizing that the equilibrium supported by most MB models is one where for each component *i*, the monotone Boolean function *f*_*i*_ will be from the largest set *U* (*b*) ⊂ *MBF* (*k*), or *L*(*b*) ⊂ *MBF* (*k*). These sets were characterized in Lemma 2.11.

#### Proof of Lemma 2.14

*Proof*. By Theorem 2.13 the models *f* = (*f*_1_, …, *f*_*N*_) that support bistability between *d* = 0 and *e* = 1 satisfy

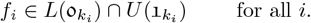

Since 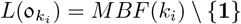 and 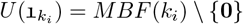 it follows that

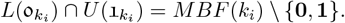

The size of this set is

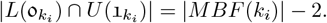

By the complementarity property in Theorem 3.6 that for any *k* and any *b* ∈ 𝔹^*k*^ the sets *U* (*b*) and *L*(*b*) satisfy

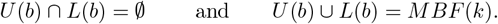

Note, however, that if *U* (*b*) = ↑ *g* and *L*(*b*) = ↓ *h* then *g* ≠↓ *h* in general, see (Table 3).

**Table 3.** Prime ideals (left) and prime filters (right) in MBF(3). Both sets are posets isomorphic to 𝔹^3^ by Birkhoff Theorem.

|  |  |  |  |  |  |
| --- | --- | --- | --- | --- | --- |
| L(000) | $L(X \vee Y \vee Z)$ | 19 | U(000) | $U(\mathbf{1})$ | 1 |
| L(010) | $L(X \vee Z)$ | 14 | U(010) | $U(Y)$ | 6 |
| L(001) | $L(X \vee Y)$ | 14 | U(001) | $U(Z)$ | 6 |
| L(100) | $L(Y \vee Z)$ | 14 | U(100) | $U(X)$ | 6 |
| L(011) | $L(X)$ | 6 | U(011) | $U(Y \wedge Z)$ | 14 |
| L(101) | $L(Y)$ | 6 | U(101) | $U(X \wedge Z)$ | 14 |
| L(110) | $L(Z)$ | 6 | U(110) | $U(X \wedge Y)$ | 14 |
| L(111) | $L(\mathbf{0})$ | 1 | U(111) | $U(X \wedge Y \wedge Z)$ | 19 |

We conclude with a result that is a direct consequence of Corollary 3.15. For illustration, see Table 3 for the table of all sets *U* (*b*), *L*(*b*) in MBF(3), which is depicted in Figure 6.

##### Lemma 3.17.

*For any b* ∈ 𝔹^*k*^ *the sets U* (*b*), *L*(*b*) ⊂ *MBF* (*k*) *satisfy*

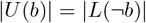

*where* ¬ *is negation*.

There are additional symmetries on the lattice *MBF* (*k*), induced by permutations *p* : {1, …, *k*} → {1, …, *k* } on the set of coordinates for vectors *b* ∈ 𝔹^*k*^. Each such permutation *p* induces isomorphism *π* : 𝔹^*k*^ → 𝔹^*k*^. Since each function *f ∈ MBF* (*k*) is uniquely determined by its truth set *T* ⊂ (*f*) 𝔹^*k*^, and the lattice operations are defined in terms of union and intersections of truth sets, the isomorphism *π* induces an automorphism

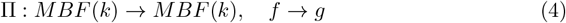

where *T* (*g*) = *π*(*T* (*f*)).

Consider collection of prime ideals 𝕃 := {*L*(*b*) *b* |∈ *MBF* (*k*)} and collection of prime filters 𝕌 := *U* (*b*) |*b* ∈ *MBF* (*k*). Each of these collections has 2^*k*^ elements, but the existence of this automorphism implies that these sets only have *k* + 1 different sizes.

##### Theorem 3.18.

*Consider collection of prime ideals* 𝕃 := {*L*(*b*) *b* ∈ *MBF* (*k*)} *and collection of prime filters* 𝕌 := {*U* (*b*) *b* ∈ *MBF* (*k*)}. *Then there is a sequence of k* + 1 *integers*

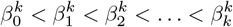

*with* 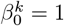, 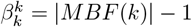, *such that*

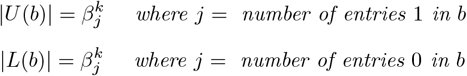

*Proof*. The fact that 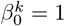, 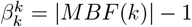 follows from Lemma 2.11.

Consider any permutation *p* : {1, …, *k*} → {1, …, *k*} on the set of coordinates for vectors *b* ∈ 𝔹^*k*^ and the induced automorphism Π in (4).

It is easy to check that Π preserves the property of join- and meet-irreducibility and therefore induces an automorphism on the set 𝕌. Since the map *π* preserves the number of 1s in the string, it follows that the size |*U* (*b*) |= |*U* (*π*(*b*)) |. Therefore, the size of |*U* (*b*) | only depends on the number of entries 1 in the string *b*.

Finally, since *MBF* (*k*) is a collection of increasing monotone Boolean functions, then *b* ≺ *c* in 𝔹^*k*^ implies *U* (*b*) ⊂ *U* (*c*) and thus *U* (*b*) ⊂ *U* (*c*). This finishes the proof for the set 𝕌. The result for the set 𝕃 follows now from Lemma 3.17.

To illustrate the Theorem recall from Table 1 that for *MBF* (2) the sizes of the sets *U* (*b*), *L*(*b*) are

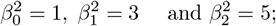

from Table 3 the sizes are

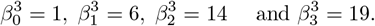

#### Proof of Theorem 2.8

*Proof*. We order nodes *x*_1_, …, *x*_*N*_. If there is no *j* such that *f*_*j*_ is decreasing in *x*_1_, then all edges with source *x*_1_ in the influence network are positive. We set *α*_1_(*x*_1_) := *x*_1_. Otherwise, if there is an *f*_*j*_ which is decreasing in *x*_1_ we set

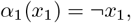

and

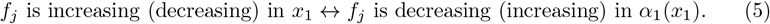

However, the update function for the node *x*_1_: *x*_1_ = *f*_1_(*x*), we need to also change this function

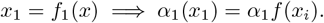

Setting *g*_1_ := *αf*_1_ it follows that

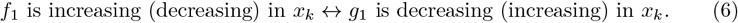

Consider now the influence network of a new model 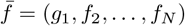 with state (*α*_1_(*x*_1_), *x*_2_, …, *x*_*N*_). By (5) all influence edges of the original network *f* that start in *x*_1_ have changed signs in the influence network of 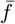; and by (6) all influence edges that terminate in *x*_1_ also changed signs.

Importantly, in any loop that passes through node *x*_1_, there were two changes, Hence, the sign of the loop did not change, and we eliminated the negative edge from *x*_1_ to *x*_*j*_ from the influence network. We conclude that the sign of every loop in the influence network of *f* is the same as in the influence network of 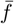.

We now consider the sources of node *x*_1_ and ask if *g*_1_ is decreasing in any of its inputs. If so, we eliminate these negative edges as described above until no negative inputs of *g*_1_ remain. We then move to the sources of *g*_1_.

Since there are no negative loops in the network *RN* when a previously considered node appears in the list of sources, the corresponding loop will have all positive edges, and no change will be required for such a node. Since there is a finite number of nodes in the network and we will only need to consider each of them for the change *α* only once, this process will terminate. Since every loop in the network RN is positive, the change of variables *α* will yield *g* with positive influence network.

## 4 Discussion

This paper presents a first step towards a mathematical understanding of the relationship between network structure and the dynamics it supports. The answer clearly depends on a class of models that is selected to represent the network dynamics; answering this question in the class of all differential equation models compatible with the network runs immediately into problems with basic concepts: What does it mean to have the same dynamics when both the class of models and the state space of every single model are uncountable?

We investigate this question within the class of monotone Boolean models where the set of models is finite (but large) and the state space is a finite set of Boolean vectors 𝔹^*N*^. In spite of this finiteness, Boolean models are very popular in systems biology as they combine the finiteness with sufficient expressiveness to capture qualitative dynamic behavior of regulatory networks.

We make two additional simplifications. First, we seek to describe the relationship between network structure and steady states, rather than more general dynamic behavior. Second, because we focus on steady states, we only consider negative loop-free (NL-free) networks. It is known [31] that negative loops are associated with the existence of attractors that consist of more than one state (for instance, complex attractors), and, in fact, the existence of such attractors requires negative loops in a network.

In networks with negative loops, some of the MBM will not support any equilibria, only complex attractors, automatically decreasing the number of MBMs supporting a given equilibrium. This is consistent with results from the toggle triad, where the prevalence/stability of single high states is lower than the corresponding prevalence in two-team networks (NL-free) networks [32]. This assumption that excludes networks with negative loops has been done for convenience; the work on understanding the connection between negative loops and complex attractors is the subject of our current work, and we believe that the approach outlined in this paper can be extended to general networks that contain negative loops.

Our results are consistent with work on the prevalence of steady states for the EMT network. Paper [33] found that positive loops are enhanced in the EMT network, which is consistent with the high prevalence of bistability in ODE simulations of EMT models. Two EMT models that are studied in this paper are NL-free, and therefore, the E and M states are the most prevalent steady states across all MBM models for these networks. Further study of the robustness of this bistability was examined in [34]. This paper studied 13 different EMT networks and showed that while these wild-type networks show the highest prevalence of E and M states in sampled ODE simulations, small perturbations of the network structure (i.e. change of edge sign) produced networks where the prevalence of E and M states decreased and the prevalence of intermediate states increased. While our results do not directly address robustness under network perturbations, our results imply that among all networks with the same number of nodes and edges, the network that has all positive edges will have the highest prevalence of the constant 0 and 1 states. Since EMT networks studied in this paper are NL-free and thus E and M states are the most prevalent, any perturbation of this network that changes the signs of the edges will result in a network with decreased prevalence of E and M states and increased prevalence of the intermediate states.

The realization that the sets of monotone Boolean functions that evaluate to either 0 or 1 as prime ideals or filters in the lattice of monotone Boolean functions with *k* inputs, allows us to prove general results that are valid for all *k*. This is significant since the size of *MBF* (*k*) grows rapidly with *k* and it is only known for *k* ≤ 9, but a construction of meet- and join-irreducible elements in *MBF* (*k*) may be possible by inductive procedure from those elements in *MBF* (*k* − 1) in a way similar to the construction of *MBF* (*k*) from *MBF* (*k* − 1) [15]. This in turn would enable explicit construction of monotone Boolean models with certain multistability properties, or,alternatively, prove that a particular network does not support multistability greater than some upper bound *q*.

Monotone Boolean function models are closely linked with ODE models with steep nonlinearities which in the limit become *Glass models* with piecewise constant nonlinearities. Each such ODE model generates a finite state transition graph (STG) that encodes its dynamics, and there is a only a finite collection of STGs that are compatible with the given network. This collection is organized in a DSGRN database which takes a form of a graph [9, 10]. Since MB network models are a subset of DSGRN collection, the work presented here should describe multi-stability in ODE network models with steep nonlinearities. The precise nature of this connection is a subject of current investigation.

*Reaction network theory* takes a different approach to modeling network dynamics using mass action kinetics. The delineation of conditions that guarantee (or preclude) multi-stability has been a subject of many papers, see for instance [35–37].

The underlying philosophy of our approach that aims to describe the full suite of network dynamics is that important dynamics will occur with higher frequency within this set. The importance of particular network dynamics is weighted by the prevalence of MBMs that support it. That is, we view a steady state that is supported by 80% of all MBMs as more important than a steady state that is only supported by 10% of all MBMs. It is certainly possible that biological control external to the network can keep the network in a (set of) MBMs which supports the dynamics with 10% prevalence, but in the absence of such information, we will assume that the more prevalent dynamics is more important.

## Acknowledgments

Part of this work has been done while the last author visited Department of Bioengineering, Indian Institute of Science, Bengaluru, India. Its hospitality is gratefully acknowledged.

